# GenomicSign: A computational method to discover unique, specific, and amplifiable signatures of target genomic sequences

**DOI:** 10.1101/2024.11.05.622192

**Authors:** S. Prasanna Kumar, Ashok Palaniappan

## Abstract

Molecular diagnostics for the rapid identification of infectious, virulent, and pathogenic organisms are key to health and global security. Such methods rely on the identification and detection of signatures possessed by the organism. In this work, we outline a computational algorithm, GenomicSign, to determine unique and amplifiable genomic signatures of a set of target sequences against a background set of non-target sequences. The set of target sequences might comprise variants of a pathogen of interest, say SARS-CoV2 virus. Unique k-mers of the consensus target sequence for a range of k-values are determined, and the threshold k-value yielding a sharp transition in the number of unique k-mers is identified as k_opt_. Corresponding unique k-mers for k ≥ k_opt_ are compared against the set of non-target sequences to identify *target-specific* unique k-mers. A pair of proximal such k-mers could enclose a potential amplicon. Primers to such pairs are designed and scored using a custom scheme to rank the potential amplicons. The top-ranked resulting amplicons are candidates for unique and amplifiable genomic signatures. The entire workflow is demonstrated using a case study with the SARS-CoV2 omicron genome. A case study distinguishing the SARS-CoV2 omicron target strain against non-target other SARS-CoV2 variants is performed to illustrate the workflow. GenomicSign has been implemented in Python and is available as an open-source software under MIT licence (https://www.github.com/apalania/GenomicSign).

## Introduction

DNA barcoding is an essential enterprise in biotechnology, biodiversity, and medicine. Emergence of novel pathogenic phenotypes as a response to selective pressures operating on infectious agents or due to zoonoses is well-documented. Mutations and horizontal gene transfer comprise mechanisms that mediate the acquisition of novel pathogenicity. Genomic barcodes of emerging agents would be useful for fingerprinting infectious strains in public health scenarios. Such barcodes might involve a genomic region unique to the agent of interest against a pool of background DNA, and could be described as genome signatures that precisely identify the agent. Precise identification of pathogenic strains is necessary for early detection, effective monitoring, and design of therapeutic interventions. It may aid environmental surveillance in the age of pandemics and constant emerging infections. Finding a unique amplifiable genomic region specific to some pathogen might comprise a valuable addition to our preparedness against infections.

Earlier methodological efforts to identify signatures of a specific population of interest against background genomes include KPATH, TOFI, TOPSI, Insignia and Neptune. However the idea of an optimal and unique k-mer co-occurring with a proximal second such k-mer yielding a candidate signature amplicon has not been explored so far. In this work, our objective is to build a pipeline called GenomicSign that facilitates the identification of such candidate genomic signatures of a given genome or set of related genomes. Central to our effort here is the notion of a ‘characteristic genome length,’ introduced in the Methods section. GenomicSign composes multiple existing bioinformatics tools to funnel out target-specific unique k-mers and then relies on an interpretation of standard primer design guidance to rank their amplifiability. GenomicSign is customizable and freely available for academic use at https://www.github.com/apalania/GenomicSign.

### Algorithm

The central premise of the algorithm is that there exists a ‘characteristic genome length’ of an organism, k_opt_, at which a dramatic increase in the number of unique (or singleton) k-mer sequences for the given genome occurs. Let n_k_ represent the number of singleton k-mers for some value of k. As k increases, n_k_ shifts from low values to high values, touching a maximum denoted by max_n_k_ at k = k_max_. For small k-values (k<k_opt_), most of the possible k-mers would be sampled *multiple times* in the genome. For k > k_max_, n_k_ smoothly decreases to a value of 1 attained when k = size of genome. Based on the above discussion, k_opt_ can be defined as the smallest k-value with n_k_ > 0.5 * max_n_k_. Essentially k_opt_ marks the transition in a two-state model of low n_k_ vs. high n_k_ values.

The first part of the algorithm consists in finding k_opt_ [Figure 1A]. The input to the algorithm includes multiple whole-genome sequences of a target organism and the set of whole-genome sequences of non-target organisms. A multiple sequence alignment of the sequences in the target set yields the consensus genome of the target organism. The consensus target sequence is then analyzed using Jellyfish to find unique k-mers for a range of k values [ref]. Parsing Jellyfish results yields a sequence of n_k_ values, whose max_n_k_ could be probed for a sharp transition at k = k_opt_. Let [k_opt_, k_max_] define a neighborhood around k_opt_. For each k in this neighborhood interval, corresponding singleton k-mers are searched against each sequence in the non-target set using cd-hit-est-2d [ref]. Those that occur in none of the genomes in the non-target set are determined. Since the genomes in the non-target set would have variable ‘characteristic lengths’, all k_opt_-mers from the target set might occur at least once in the non-target set in some genome. By scanning singleton k-mers in the neighborhood interval of k_opt_, we would eventually identify the k-value at which singleton k-mers from the target set are not found in the non-target set at all. Singleton k-mers in the target set for the least such k-value comprise seed choices for the design of genomic signatures, and are called candidate target-specific unique (CTU) sequences.

**Figure 1.**
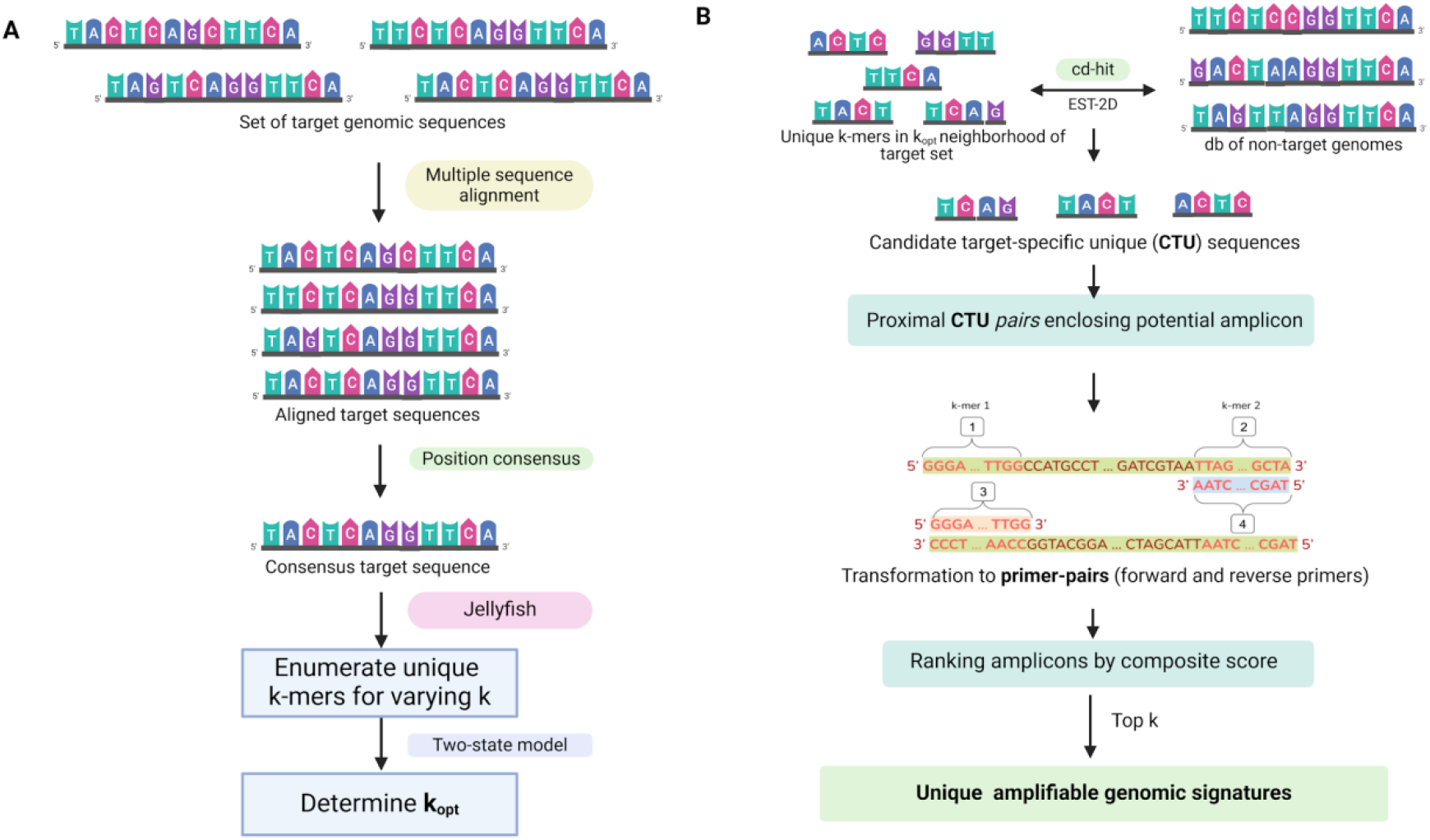
GenomicSign algorithm, graphical representation. **(A)** Working with the target set of sequences to arrive at k_opt_, the transition point between low and high n_k_ values; (**B)** Using the unique k-mers from **A** to identify CTU sequences and rank amplicons.

To design an amplicon, we will need a pair of CTU sequences within a reasonable genomic interval of each other. In general amplicons need to span a genomic interval of ∼80-200 bp for polymerase chain reactions (PCR). Thus pairs of CTU sequences that occur in proximity to span a specific genomic interval are identified as potential amplicons of interest. Addition of nucleotides to a unique k-mer yields a longer unique sequence clearly, implying that the amplicons thus circumscribed could be candidate genomic signatures. Primer pairs are obtained from such CTU sequence pairs (Figure 1B). Each primer is then ranked using the scoring scheme presented in Table 1. A ViennaRNA utility called RNAcofold (with ‘dna’ option) is used to detect annealing between the forward and reverse primers [ref]. The magnitude of dimerization is quantified by the nesting level in the dot bracket representation. Intraprimer annealing might manifest as multi-branched junctions. GenomicSign uses Schudoma’s mdg_dt.py [ref] to quantify multiloops within a primer. The putative amplicons are ranked based on the cumulative score of the forward and reverse primers and the most promising candidate target-specific genomic signatures are returned.

**Table 1.**
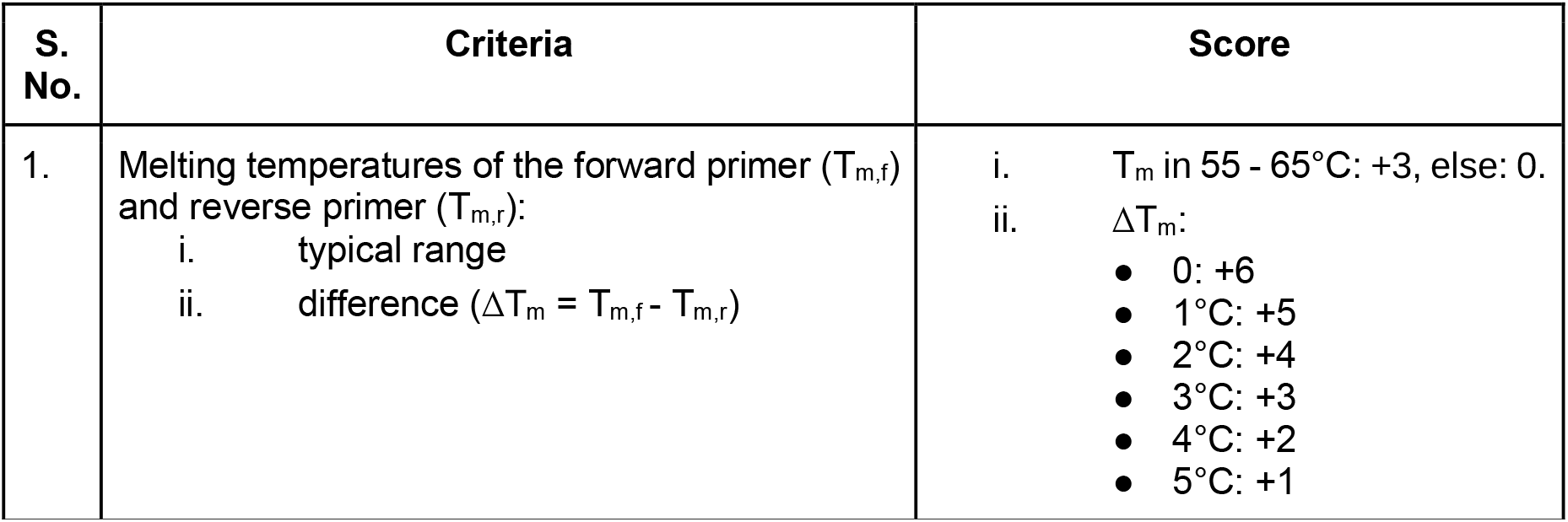
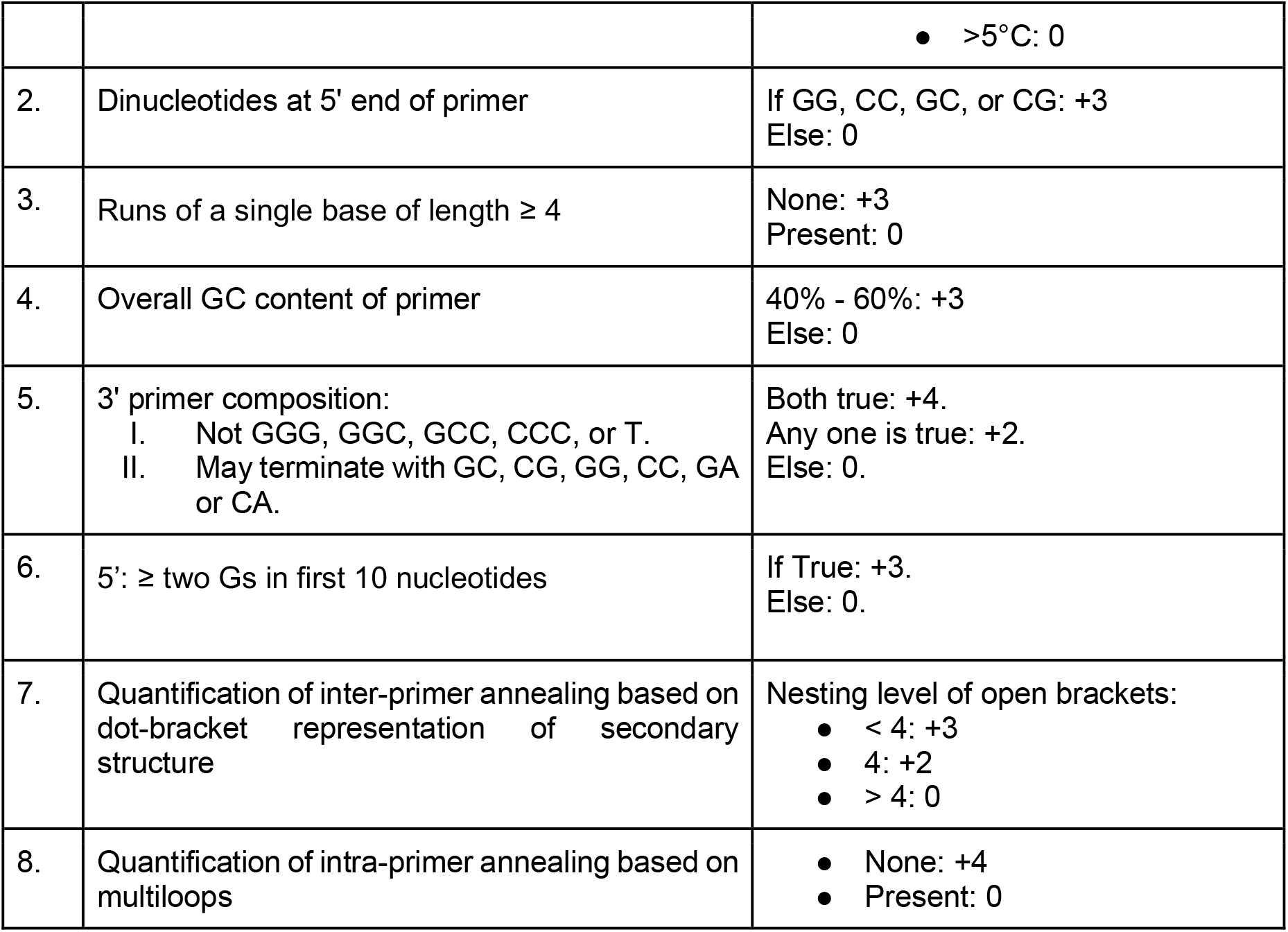
A customizable scoring scheme for primer design. ViennaRNA was used to to quantify inter-primer annealing based on the nesting level of dot-bracket representation.Satisfaction of constraints is rewarded.

| S. No. | Criteria | Score |
| --- | --- | --- |
| 1. | Melting temperatures of the forward primer ($T_{m,f}$) and reverse primer ($T_{m,r}$):<br>i. typical range<br>ii. difference ($\Delta T_m = T_{m,f} - T_{m,r}$) | i. $T_m$ in 55 - 65°C: +3, else: 0.<br>ii. $\Delta T_m$:<br>• 0: +6<br>• 1°C: +5<br>• 2°C: +4<br>• 3°C: +3<br>• 4°C: +2<br>• 5°C: +1 |
|  |  | • $>5^{\circ}\text{C}$: 0 |
| 2. | Dinucleotides at 5' end of primer | If GG, CC, GC, or CG: +3<br>Else: 0 |
| 3. | Runs of a single base of length $\geq 4$ | None: +3<br>Present: 0 |
| 4. | Overall GC content of primer | 40% - 60%: +3<br>Else: 0 |
| 5. | 3' primer composition:<br>I. Not GGG, GGC, GCC, CCC, or T.<br>II. May terminate with GC, CG, GG, CC, GA or CA. | Both true: +4.<br>Any one is true: +2.<br>Else: 0. |
| 6. | 5': ≥ two Gs in first 10 nucleotides | If True: +3.<br>Else: 0. |
| 7. | Quantification of inter-primer annealing based on dot-bracket representation of secondary structure | Nesting level of open brackets:<br>• $< 4$: +3<br>• 4: +2<br>• $> 4$: 0 |
| 8. | Quantification of intra-primer annealing based on multiloops | • None: +4<br>• Present: 0 |

### Case study

To demonstrate GenomicSign, we performed a case study to identify genomic signatures of SARS-CoV-2 (hCoV-19) virus specific to a variant of concern (VOC). In particular we sought unique signatures of the omicron VOC against other serotypes of SARS-CoV-2. Omicron VOC (29903 bp) is the predominant infectious hCoV-19 strain circulating in the world today. To build datasets of target and non-target genomes, we used the Global Initiative on Sharing All Influenza Data (GISAID) [ref.] to search for multiple SARS-CoV-2 VOCs, which are named by the geographical distribution of first occurrence. The target dataset comprised 10 Omicron VOC genomes (Hong Kong). The non-target dataset comprised of 10 genomes each of seven other VOCs, namely Alpha (UK), Beta (Africa), Gamma (Japan), Delta (India), Lambda (Peru), Mu (Colombia), and VUM GH (France), for a total of 70 non-target genomes. Table 2 shows the accession IDs of the genomes used in the study; the fasta sequences are provided in the GitHub repository.

**Table 2.** VOCs used in the study and their unique GISAID accession IDs. Ten genomes were used to represent each VOC.

| S. No. | Variant | Target/Non-Target | Accession IDs |
| --- | --- | --- | --- |
| 1. | Omicron | Target | EPI_ISL_12417949, EPI_ISL_12417952, PI_ISL_12417953, EPI_ISL_12417962, EPI_ISL_12417960, EPI_ISL_12417959, EPI_ISL_12417958, EPI_ISL_12417957, EPI_ISL_12417956, EPI_ISL_12417954 |
| 2. | Alpha | Non-target | EPI_ISL_11059129, EPI_ISL_11059128, EPI_ISL_11059113, EPI_ISL_11059059, EPI_ISL_11059037, EPI_ISL_10993510, EPI_ISL_10993509, EPI_ISL_10993505, EPI_ISL_10993503, EPI_ISL_10993502 |
| 3. | Beta | Non-target | EPI_ISL_13370871, EPI_ISL_13370868, EPI_ISL_13370815, EPI_ISL_13183619, EPI_ISL_13183618, EPI_ISL_13183607, EPI_ISL_13183580, EPI_ISL_13037944, EPI_ISL_13037942, EPI_ISL_13037938 |
| 4. | Gamma | Non-target | EPI_ISL_6228367, EPI_ISL_5716982, EPI_ISL_5461734, EPI_ISL_4720113, EPI_ISL_4713840, EPI_ISL_4713792, EPI_ISL_4713749, EPI_ISL_4703968, EPI_ISL_3983486, EPI_ISL_3543154 |
| 5. | Delta | Non-target | EPI_ISL_13408888, EPI_ISL_13408887, EPI_ISL_13408885, EPI_ISL_13390372, EPI_ISL_13390371, EPI_ISL_13390369, EPI_ISL_13390368, EPI_ISL_13390366, EPI_ISL_13390365, EPI_ISL_13390364 |
| 6. | Lambda | Non-target | EPI_ISL_13107634, EPI_ISL_12934522, EPI_ISL_11840897, EPI_ISL_10328969, EPI_ISL_8882505, EPI_ISL_8874883, EPI_ISL_8797236, EPI_ISL_8797233, EPI_ISL_8797232, EPI_ISL_8797231 |
| 7. | Mu | Non-target | EPI_ISL_8844686, EPI_ISL_8844685, EPI_ISL_8844683, EPI_ISL_8844682, EPI_ISL_8844681, EPI_ISL_8844680, EPI_ISL_8844679, EPI_ISL_8844678, EPI_ISL_8844677, EPI_ISL_8844676 |
| 8. | VUM GH | Non-target | EPI_ISL_11580635, EPI_ISL_11580559, EPI_ISL_11225839, EPI_ISL_11036995, EPI_ISL_11036903, EPI_ISL_11036869, EPI_ISL_11036865, EPI_ISL_11036862, EPI_ISL_11036860, EPI_ISL_10947040 |

To identify potential genomic signatures specific to the Omicron variant (and absent in the other VOCs), the GenomicSign pipeline is executed. Following alignment of the target set of sequences using Clustal Omega with default parameters [ref], the target consensus was determined using Biopython [ref]. Calling Jellyfish on the consensus sequence and subsequent analysis yielded k_max_ = 17 and k_opt_ = 9 [Figure 2A]. In the next step, cd-hit-est-2d was used to search the unique k-mers for k ≥ k_opt_ against the consolidated set of non-target VOC genomes (parameters: identity = 1.0, word size = 8). Strikingly, none of the unique k-mers at k=k_opt_ and k=k_opt_+1 yielded specificity to the target. Scanning with singleton k-mers at *k = 11* yielded 11,235 CTU sequences [Figure 2B]. Searching for proximal sequences from this set yielded just one pair of CTU sequences, enclosing an 81-nt potential amplicon. The CTU sequences in this pair were extended to the desired primer length (20 oligo-nt), to yield a cumulative score ∼ 38. Table 3 presents a summary of the findings.

**Table 3.** GenomicSign pipeline applied to COVID VOCs. Summary of the intermediate results upon sequential execution of GenomiSign on Omicron target genomes against other non-target COVID VOCs. The pipeline culminates in the identification of a single 81-bp amplicon.

|  | Pipeline stage | Result |
| --- | --- | --- |
| 1. | Input: 10 hCoV Omicron VOC genomes | Target consensus sequence |
| 2. | Jellyfish: find number of unique k-mers for k in range (4, 20) and determine characteristic length, $k_{opt}$ | $k_{max} = 17$, midpoint $\sim 15,000$<br>$\Rightarrow k_{opt} = 9$ |
| 3. | Cd-hit-est-2d: find target-specific unique k-mers for k in $[k_{opt}, k_{max}]$ neighbourhood | CTU: 11,235 11-mers |
| 4. | Proximal CTU pairs | One 81-nt potential amplicon with genomic coordinates: (272, 353) |
| 5. | Primer scoring | Cumulative score of identified potential amplicon = 38 |

**Figure 2.**
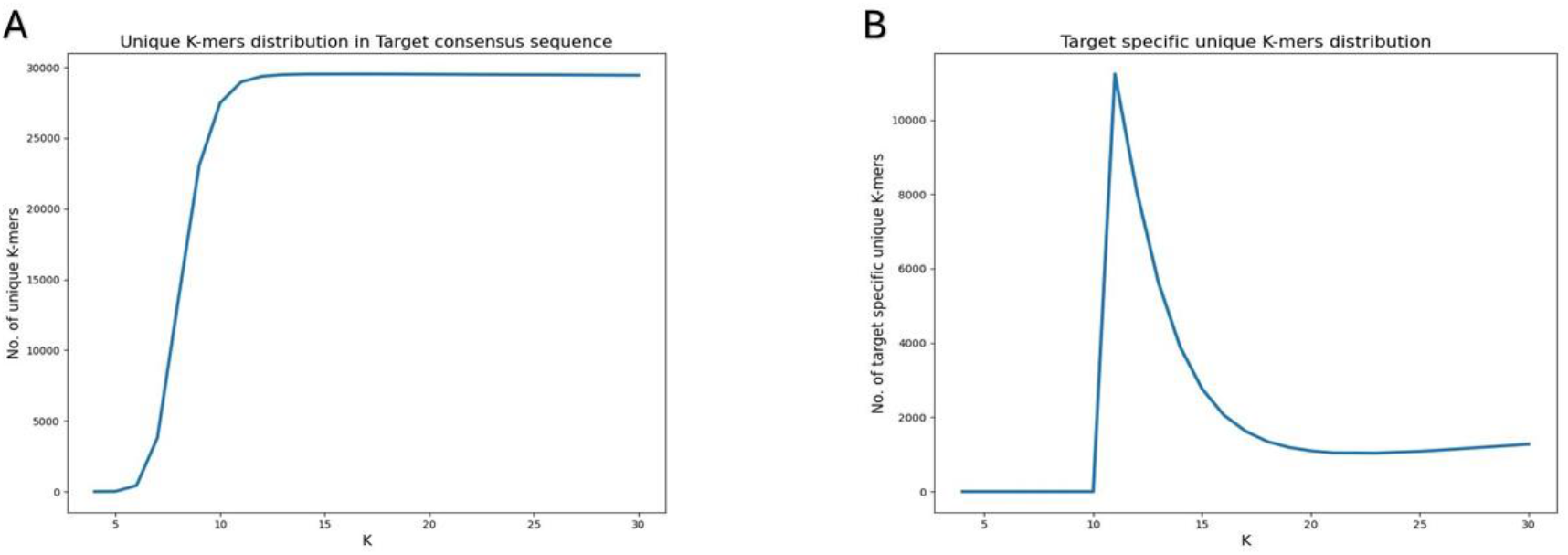
Identification of k_opt_ and CTU sequences. **(A)** Distribution of unique k-mers depicted as n_k_ vs k from Jellyfish results for the target set of the case study. A two-state model of high vs low n_k_ values can be seen, and the transition occurs at k_opt_ = 9; **(B)** Distribution of CTU sequences depicted as #CTU sequences vs k for varying values of k, obtained using cd-hit-est-2d. The low values can be attributed to low-complexity and conserved k-mers for k < 11. The spike is seen at k = 11, yielding 11,235 CTU sequences. Changing the word size does not affect the results.

The identified 81 bp amplicon lies in the coding region of the ORF1ab gene, just after the 5’ start [Figure 3; ref]. It is necessary to validate whether the said final signature contained in the amplicon does identify the target genomes of interest precisely. Toward this, we executed BLAST search of the amplicon signature sequence against the GISAID database (with default parameters; ref). Unsurprisingly, all the top hits in the search were marked as Omicron VOC genomes. Variants BA.2.86.1 and JN.2 are also returned, both referring to Omicron variants in PANGO nomenclature [ref]. An analysis of the top *30* hits shows ungapped full-length alignments with 100% identity and E-value ∼ 1e-34, starting ∼ 271±36 in the sequences. These results may be viewed against the fact that the consensus of the target set is not identical to any of the sequences in the target set. The query is specific against non-target hCoV VOC genomes, with the only false positive being a Delta variant [ref]. Given that the query amplicon does not occur verbatim in any of the target genomes, the above observations provide evidence for the discovery of a signature. The complete results are included in Supplementary Information.

**Figure 3.**
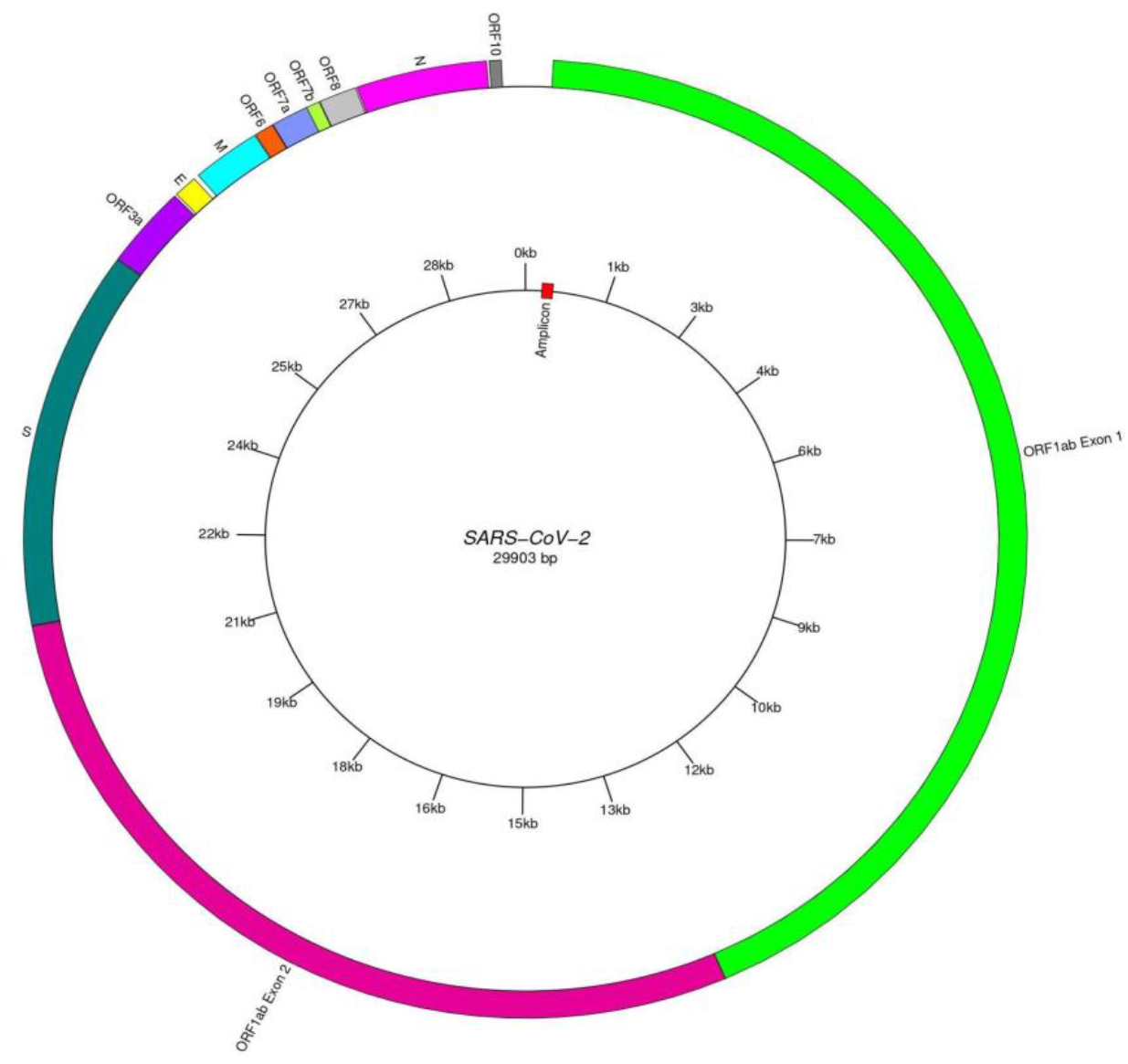
Representation of the SARS-CoV-2 Omicron genome annotated with genes and the locus of the 81-nt ‘amplicon’ identified using GenomicSign. The potential signature lies within the Exon 1 of the ORF1ab gene. No other signatures were identified by GenomicSign for the case study considered.

### Implementation

GenomicSign has been implemented in Python3.x with third-party requirements for multiple sequence alignment, RNA hybridization, and multiloop detection. Customizable parameters (with their default values) are given below:

i Lower (4) and upper bounds (30) for the range of k-values explored in enumerating unique k-mers.
ii Lower (80) and upper (200) bounds for the interval size necessary to define proximal genomic signatures; suggested to fall within 40-400 bases. GenomicSign scans for CTU sequences separated from each other at a genomic distance in this range. The corresponding pairs together with the interval sequence constitute a potential amplicon.
iii Primer length (20); can be set from 18-30 oligonucleotides to ensure optimal primer function.

The elastic and modular design of GenomicSign facilitate the reuse of its components for various other tasks; for e.g, the pipeline could be adapted to find a set of k-mers unique to the set of target genomes and/or specific against non-target background DNA. It could work with sets of genomes or alternatively libraries of sequence fragments; all with or without background discrimination. It could be used to merely rank a given set of potential amplicons, skipping the upstream analysis. The practical use of genomic signatures for precise and rapid genotyping is dependent on the existence of amplicons. By requiring two, not one, unique and specific k-mers to be within an amplicon distance of each other, GenomicSign engenders a technique for identifying promising potential amplicons.

### Limitations

The case study suggests that GenomicSign would be able to yield representative amplicons unique to target genomes. GenomicSign is not entirely deterministic (since multiple sequence alignment is an NP-hard problem). Given the existence of uncertainties that inhere with the use of heuristics, theoretically identified amplicons require experimental validation for proof and further clinical utility. Divergent target genomes that resist a useful consensus sequence definition might yield amplicons that are not realistic of the actual target genomes. Future studies may address finding a workaround to these limitations. It is suggested to work with a well-defined target set (or single sequences) and validate the findings in a clinical context prior to deployment in diagnostic assays.

## Conclusions

GenomicSign algorithm has been conceived to recognize genomic signatures of a target organism against some background. It has been implemented as a simple and efficient pipeline to zero in on and rank proximal genomic signatures representing potential amplicons. Robust testing may be necessary in the future on the roadmap to precision diagnosis and surveillance of biological agents.

## Acknowledgements

We would like to thank SASTRA Deemed University for infrastructure and support. We gratefully acknowledge all data contributors, i.e., the Authors and their Originating laboratories responsible for obtaining the specimens, and their Submitting laboratories for generating the genetic sequence and metadata and sharing via the GISAID Initiative, on which the research case study here is based. AP would like to acknowledge support from BT/PR40144/BTIS/137/46/2022 and BT/PR40150/BTIS/137/81/2023.

## Notes

### Competing Interest Statement

The authors have declared no competing interest.

## References

Bae, Jeongmin, et al. “GPrimer: A Fast GPU-Based Pipeline for Primer Design for qPCR Experiments.” BMC Bioinformatics, vol. 22, no. 1, Apr. 2021, p. 220. PubMed, 10.1186/s12859-021-04133-4.

Bohlin, Jon. “Genomic Signatures in Microbes -- Properties and Applications.” TheScientificWorldJournal, vol. 11, Mar. 2011, pp. 715–25. PubMed, 10.1100/tsw.2011.70.

Campbell, A., et al. “Genome Signature Comparisons among Prokaryote, Plasmid, and Mitochondrial DNA.” Proceedings of the National Academy of Sciences of the United States of America, vol. 96, no. 16, Aug. 1999, pp. 9184–89. PubMed, 10.1073/pnas.96.16.9184.

Cock, Peter J. A., et al. “Biopython: Freely Available Python Tools for Computational Molecular Biology and Bioinformatics.” Bioinformatics, vol. 25, no. 11, June 2009, pp. 1422–23. (Crossref), 10.1093/bioinformatics/btp163.

Conant, Gavin C., and Kenneth H. Wolfe. “GenomeVx: Simple Web-Based Creation of Editable Circular Chromosome Maps.” Bioinformatics, vol. 24, no. 6, Mar. 2008, pp. 861–62. (Crossref), 10.1093/bioinformatics/btm598.

David, Benjamin M., et al. “A Reinforcement Learning Framework for Pooled Oligonucleotide Design.” Bioinformatics (Oxford, England), vol. 38, no. 8, Apr. 2022, pp. 2219–25. PubMed, 10.1093/bioinformatics/btac073.

Deiman, Birgit, et al. “Characteristics and Applications of Nucleic Acid Sequence-Based Amplification (NASBA).” Molecular Biotechnology, vol. 20, no. 2, 2002, pp. 163–80. (Crossref), 10.1385/MB:20:2:163.

Dwivedi, Bhakti, et al. “PhiSiGns: An Online Tool to Identify Signature Genes in Phages and Design PCR Primers for Examining Phage Diversity.” BMC Bioinformatics, vol. 13, no. 1, Dec. 2012, p. 37. (Crossref), 10.1186/1471-2105-13-37.

Dyer, Betsey Dexter, et al. “Classification and Regression Tree (CART) Analyses of Genomic Signatures Reveal Sets of Tetramers That Discriminate Temperature Optima of Archaea and Bacteria.” Archaea, vol. 2, no. 3, Jan. 2008, pp. 159–67. (Crossref), 10.1155/2008/829730.

Fu, Limin, et al. “CD-HIT: Accelerated for Clustering the next-Generation Sequencing Data.” Bioinformatics, vol. 28, no. 23, Dec. 2012, pp. 3150–52. (Crossref), 10.1093/bioinformatics/bts565.

Hafez, Ahmed, et al. “SeqEditor: An Application for Primer Design and Sequence Analysis with or without GTF/GFF Files.” Bioinformatics (Oxford, England), vol. 37, no. 11, July 2021, pp. 1610–12. PubMed, 10.1093/bioinformatics/btaa903.

Jernigan, Robert W., and Robert H. Baran. “Pervasive Properties of the Genomic Signature.” BMC Genomics, vol. 3, no. 1, Aug. 2002, p. 23. 10.1186/1471-2164-3-23.

Kalendar, Ruslan. “A Guide to Using FASTPCR Software for PCR, In Silico PCR, and Oligonucleotide Analysis.” Methods in Molecular Biology (Clifton, N.J.), vol. 2392, 2022, pp. 223–43. PubMed, 10.1007/978-1-0716-1799-1_16.

Karimi, Ramin, and Andras Hajdu. “HTSFinder: Powerful Pipeline of DNA Signature Discovery by Parallel and Distributed Computing.” Evolutionary Bioinformatics, vol. 12, Jan. 2016, p. EBO.S35545. (Crossref), 10.4137/EBO.S35545.

Karlin, S., and C. Burge. “Dinucleotide Relative Abundance Extremes: A Genomic Signature.” Trends in Genetics: TIG, vol. 11, no. 7, July 1995, pp. 283–90. PubMed, 10.1016/s0168-9525(00)89076-9.

Liu, Jia, et al. “MetaFunPrimer: An Environment-Specific, High-Throughput Primer Design Tool for Improved Quantification of Target Genes.” mSystems, vol. 6, no. 5, Oct. 2021, p. e0020121. PubMed, 10.1128/mSystems.00201-21.

Lopez-Rincon, Alejandro, et al. “Classification and Specific Primer Design for Accurate Detection of SARS-CoV-2 Using Deep Learning.” Scientific Reports, vol. 11, no. 1, Jan. 2021, p. 947. (Crossref), 10.1038/s41598-020-80363-5.

Lorenz, Ronny, et al. “ViennaRNA Package 2.0.” Algorithms for Molecular Biology, vol. 6, no. 1, Dec. 2011, p. 26. (Crossref), 10.1186/1748-7188-6-26.

Marçais, Guillaume, and Carl Kingsford. “A Fast, Lock-Free Approach for Efficient Parallel Counting of Occurrences of k-Mers.” Bioinformatics, vol. 27, no. 6, Mar. 2011, pp. 764–70. (Crossref), 10.1093/bioinformatics/btr011.

Marinier, Eric, et al. “Neptune: A Bioinformatics Tool for Rapid Discovery of Genomic Variation in Bacterial Populations.” Nucleic Acids Research, vol. 45, no. 18, Oct. 2017, pp. e159–e159. (Crossref), 10.1093/nar/gkx702.

O’Toole, Áine, et al. “Pango Lineage Designation and Assignment Using SARS-CoV-2 Spike Gene Nucleotide Sequences.” BMC Genomics, vol. 23, no. 1, Dec. 2022, p. 121. (Crossref), 10.1186/s12864-022-08358-2.

Pertsemlidis, Alexander, and John W. Fondon. “Having a BLAST with Bioinformatics (and Avoiding BLASTphemy).” Genome Biology, vol. 2, no. 10, Oct. 2001, pp. 1–10. genomebiology.biomedcentral.com, 10.1186/gb-2001-2-10-reviews2002.

Phillippy, A. M., et al. “Insignia: A DNA Signature Search Web Server for Diagnostic Assay Development.” Nucleic Acids Research, vol. 37, no. Web Server, July 2009, pp. W229–34. (Crossref), 10.1093/nar/gkp286.

Podell, Sheila, and Terry Gaasterland. “DarkHorse: A Method for Genome-Wide Prediction of Horizontal Gene Transfer.” Genome Biology, vol. 8, no. 2, 2007, p. R16. PubMed, 10.1186/gb-2007-8-2-r16.

Pride, David T., and Thomas Schoenfeld. “Genome Signature Analysis of Thermal Virus Metagenomes Reveals Archaea and Thermophilic Signatures.” BMC Genomics, vol. 9, no. 1, 2008, p. 420. (Crossref), 10.1186/1471-2164-9-420.

Purohit, Hj, et al. “Identification of Signature and Primers Specific to Genus Pseudomonas Using Mismatched Patterns of 16S rDNA Sequences.” BMC Bioinformatics, vol. 4, no. 1, May 2003, p. 19. (Crossref), 10.1186/1471-2105-4-19.

Rambaut, Andrew, et al. “A Dynamic Nomenclature Proposal for SARS-CoV-2 Lineages to Assist Genomic Epidemiology.” Nature Microbiology, vol. 5, no. 11, July 2020, pp. 1403–07. (Crossref), 10.1038/s41564-020-0770-5.

Rose, T. “CODEHOP (COnsensus-DEgenerate Hybrid Oligonucleotide Primer) PCR Primer Design.” Nucleic Acids Research, vol. 31, no. 13, July 2003, pp. 3763–66. (Crossref), 10.1093/nar/gkg524.

Rose, T. M., et al. “Consensus-Degenerate Hybrid Oligonucleotide Primers for Amplification of Distantly Related Sequences.” Nucleic Acids Research, vol. 26, no. 7, Apr. 1998, pp. 1628–35. (Crossref), 10.1093/nar/26.7.1628.

Schönung, Maximilian, et al. “AmpliconDesign - an Interactive Web Server for the Design of High-Throughput Targeted DNA Methylation Assays.” Epigenetics, vol. 16, no. 9, Sept. 2021, pp. 933–39. PubMed, 10.1080/15592294.2020.1834921.

Schudoma, Christian, et al. “Sequence–Structure Relationships in RNA Loops: Establishing the Basis for Loop Homology Modeling.” Nucleic Acids Research, vol. 38, no. 3, Jan. 2010, pp. 970–80. (Crossref), 10.1093/nar/gkp1010.

Shirshikov, Fedor V., et al. “MorphoCatcher: A Multiple-Alignment Based Web Tool for Target Selection and Designing Taxon-Specific Primers in the Loop-Mediated Isothermal Amplification Method.” PeerJ, vol. 7, Apr. 2019, p. e6801. (Crossref), 10.7717/peerj.6801.

Shu, Yuelong, and John McCauley. “GISAID: Global Initiative on Sharing All Influenza Data – from Vision to Reality.” Eurosurveillance, vol. 22, no. 13, Mar. 2017. (Crossref), 10.2807/1560-7917.ES.2017.22.13.30494.

Sievers, Fabian, et al. “Fast, Scalable Generation of High-Quality Protein Multiple Sequence Alignments Using Clustal Omega.” Molecular Systems Biology, vol. 7, Oct. 2011, p. 539. PubMed, 10.1038/msb.2011.75.

Slezak, T. “Comparative Genomics Tools Applied to Bioterrorism Defence.” Briefings in Bioinformatics, vol. 4, no. 2, Jan. 2003, pp. 133–49. 10.1093/bib/4.2.133.

Touati, Rabeb, et al. “Comparative Genomic Signature Representations of the Emerging COVID-19 Coronavirus and Other Coronaviruses: High Identity and Possible Recombination between Bat and Pangolin Coronaviruses.” Genomics, vol. 112, no. 6, Nov. 2020, pp. 4189–202. 10.1016/j.ygeno.2020.07.003.

Tuteja, Amit, et al. “GSIT: An Integrated Web-Tool for Identification of Genomic Signatures in Highly Similar DNA Sequences.” Bioinformation, vol. 10, no. 8, 2014, pp. 551–54. PubMed, 10.6026/97320630010551.

Varliero, Gilda, et al. “PhyloPrimer: A Taxon-Specific Oligonucleotide Design Platform.” PeerJ, vol. 9, Apr. 2021, p. e11120. (Crossref), 10.7717/peerj.11120.

Vijaya Satya Ravi, Nela Zavaljevski, et al. “A High-Throughput Pipeline for Designing Microarray-Based Pathogen Diagnostic Assays.” BMC Bioinformatics, vol. 9, no. 1, Dec. 2008, p. 185. (Crossref), 10.1186/1471-2105-9-185.

Vijaya Satya Ravi, Kamal Kumar, et al. “A High-Throughput Pipeline for the Design of Real-Time PCR Signatures.” BMC Bioinformatics, vol. 11, no. 1, Dec. 2010, p. 340. (Crossref), 10.1186/1471-2105-11-340.

Vitiello, Antonio, et al. “Advances in the Omicron Variant Development.” Journal of Internal Medicine, vol. 292, no. 1, July 2022, pp. 81–90. PubMed, 10.1111/joim.13478.

Wright, Erik S., and Kalin H. Vetsigian. “DesignSignatures: A Tool for Designing Primers That Yields Amplicons with Distinct Signatures.” Bioinformatics (Oxford, England), vol. 32, no. 10, May 2016, pp. 1565–67. PubMed, 10.1093/bioinformatics/btw047.

Ye, Kai, et al. “Mining Unique-m Substrings from Genomes.” Journal of Proteomics & Bioinformatics, vol. 03, no. 03, 2010, pp. 099–103. 10.4172/jpb.1000127.

